# Macrophage Migration Inhibitory Factor on Apoptotic Extracellular Vesicles Regulates Compensatory Proliferation

**DOI:** 10.1101/2023.06.14.544889

**Authors:** Safia A. Essien, Ivanshi Ahuja, George T. Eisenhoffer

## Abstract

Apoptotic cells can signal to neighboring cells to stimulate proliferation and compensate for cell loss to maintain tissue homeostasis. While apoptotic cell-derived extracellular vesicles (AEVs) can transmit instructional cues to mediate communication with neighboring cells, the molecular mechanisms that induce cell division are not well understood. Here we show that macrophage migration inhibitory factor (MIF)-containing AEVs regulate compensatory proliferation via ERK signaling in epithelial stem cells of larval zebrafish. Time-lapse imaging showed efferocytosis of AEVs from dying epithelial stem cells by healthy neighboring stem cells. Proteomic and ultrastructure analysis of purified AEVs identified MIF localization on the AEV surface. Pharmacological inhibition or genetic mutation of MIF, or its cognate receptor CD74, decreased levels of phosphorylated ERK and compensatory proliferation in the neighboring epithelial stem cells. Disruption of MIF activity also caused decreased numbers of macrophages patrolling near AEVs, while depletion of the macrophage lineage resulted in a reduced proliferative response by the epithelial stem cells. We propose that AEVs carrying MIF directly stimulate epithelial stem cell repopulation and guide macrophages to cell non-autonomously induce localized proliferation to sustain overall cell numbers during tissue maintenance.

## INTRODUCTION

The ability to maintain epithelial tissue homeostasis has important implications for the health of multi-cellular organisms. Failure to adequately replace dead or missing cells can predispose epithelia to failed tissue maintenance, loss of barrier function^1–3^, and increased susceptibility to infection^4^. Alternatively, unchecked cell growth with minimal removal of defective cells can result in hyperplasia^5^, a hallmark of cancer. Studies in *Drosophila* have shown that dying cells are able to secrete mitogenic signals to neighboring cells to stimulate proliferation^6^. Similar effects were observed in Hydra through the transfer of Wnt3 from dying cells to facilitate regeneration^7^. While the link between dying cells and proliferation has long been established, what is not entirely clear is how signals are transmitted from the dying cell to neighboring cells. Recent work has proposed apoptotic bodies or apoptotic extracellular vesicles as transporters of mitogenic signals from dying cells to neighboring cells ^8–10^.

Cells undergoing programmed cell death fragment into membrane bound vesicles that are approximately 1-5µm in diameter ^11–13^, called apoptotic bodies^11^ or apoptotic extracellular vesicles (AEVs), to prevent the contents from spilling out into the extracellular space. Similar to other extracellular vesicles, AEVs are enriched in contents that can regulate or interact with neighboring cells. For instance, AEVs can participate in the horizontal transfer of DNA^14,15^, microRNA^16^, splicing factors^17^, and biologically active proteins^18^. While apoptosis has been dogmatically regarded as an “immunologically silent” form of cell death eliciting minimal inflammation compared to necrosis^19^, damage associated molecular patterns (DAMPs) such as Hsp70 and HMGB1 have been observed in AEVs^20^. Yet, how the contents of AEVs may regulate the local microenvironment after cell death in living tissues remains poorly understood.

The technical challenges to perturb living epithelia in the presence of intact immune system and image subsequent changes in real time has thus far prevented a detailed characterization of how apoptosis can stimulate proliferation. Zebrafish larvae possess an experimentally accessible bi-layered epidermis that is similar in structure and function to those coating organ systems in mammals^21–23^, providing a system to rapidly interrogate the coordination of apoptosis and proliferation. The keratinocytes in the basal layer serve as the resident stem cell population that contributes to all of the strata in the adult epidermis^24^, and also express defined markers found in epithelial stem cells such as TP63^25–27^. Our previous work showed that these basal stem cells contribute to the clearance of AEVs that stimulate their proliferation in the tail fin epidermis of zebrafish larvae^9^. Here we performed proteomic analysis of purified AEVs in zebrafish and identified proteins associated with tissue regeneration and modulation of the immune system. We further characterized Macrophage Migration Inhibitory Factor (MIF) as a putative regulator of AEV-mediated signaling during epithelial tissue maintenance. MIF has been characterized as a cytokine, chemokine, and molecular chaperone^28^, and despite its name, plays a role in leukocyte recruitment^29–31^ and proliferation and migration of epithelial cells^32^. These data suggest MIF is capable of exerting both mitogenic and immunogenic effects, yet how these are regulated during apoptosis and compensatory proliferation *in vivo* are not well understood.

This study investigates the role of MIF in apoptosis-induced proliferation of the basal epithelial stem cells in zebrafish larvae. We show that apoptotic extracellular vesicles (AEVs) have MIF on their surface, which stimulates proliferation in surrounding epithelial stem cells via the upregulation of phosphorylated ERK. Moreover, our findings indicate that apoptosis stimulates increased mobilization of macrophages, but their contribution to AEV engulfment and clearance is minimal. Our data suggest that AEVs carrying MIF stimulate macrophages to participate in compensatory proliferation in a cell non-autonomous manner. Together, these findings highlight the dynamic interplay between AEVs, neighboring epithelial stem cells, and macrophages during resolution of cell death and maintenance of overall cell numbers.

## RESULTS

### Proteomic analysis of epithelial stem cell derived-AEVs (esAEVs) identifies proteins associated with wound healing and regeneration

We used a zebrafish model to induce death in a subset of the basal stem cells in the bi-layered larval epidermis ^21,33^. The *zc1036* GAL4 enhancer trap (BASAL-GET) line was used to drive mosaic expression of the bacterial enzyme *nsfB*, or nitroreductase (NTR) fused to mCherry ^34^ in basal epithelial stem cells^21^ (Figure 1A). After the addition of Metronidazole (referred to as MTZ) to 4-day post-fertilization (dpf) larvae for 4 hours (Figure 1B), the NTR-positive cells convert MTZ into a cytotoxic byproduct that results in DNA damage^33^ and apoptosis^35^. The dying epithelial stem cells display classic markers of apoptosis, such as increased activated-caspase 3 (Figure 1C), and the formation of apoptotic extracellular vesicles (AEVs) *in vivo* (Figure 1D, Supp. Movie 1). The dynamics of epithelial stem cell-derived AEV (esAEV) biogenesis was captured using time-lapse confocal imaging. Formation of esAEVs was observed within one hour of cell shrinkage and had an average diameter of 2.41 microns (Figure 1E). Due to the mosaic nature of our genetic system (Figure 1F’), we can visualize the interaction of dying epithelial stem cells with the remaining healthy neighbor cells (Figure 1F’’). Between 8 −18 hours post-treatment (hpt) with MTZ, TP-63 positive epithelial stem cells were visualized engulfing AEVs and apoptotic cell corpses (Figure 1F’’’, Supp. Movie 2). The most engulfment events were observed at 8 hpt (Figure 1G), with an average of 15 individual epithelial stem cells engulfing esAEVs, and decreased over time. By 18hpt, we observed a significant increase in the number of actively proliferating cells that incorporated 5-bromo-2’-deoxyuridine (BrdU) (Figure 1H). These findings are in line with our previous studies that demonstrated that ∼63% of engulfing epithelial stem cells went on to divide ^9^ and support the mitogenic potential of esAEVs.

**Figure 1.**
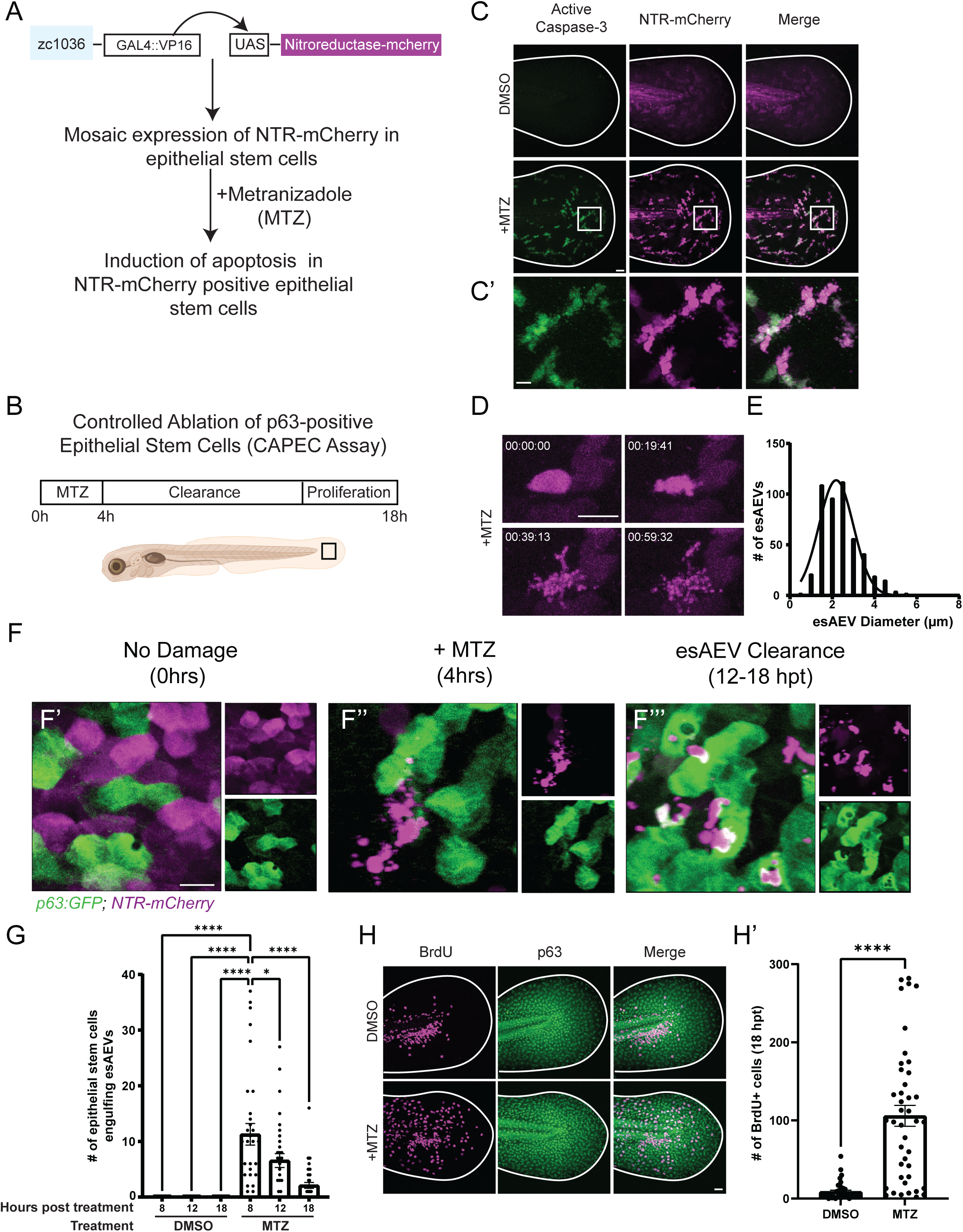
Characterization of esAEV biogenesis. (A) A description of the *Et(Gal4-VP16)^zc^*^1036^*^A^, Tg(UAS-1b:nsfB-mCherry)^c^*^264^ transgenic line. This study utilizes a GAL4/UAS zebrafish line to express the bacterial enzyme nitroreductase fused to mCherry (NTR-mCherry) in a subset of epithelial stem cells. The addition of 10mM Metronidazole (MTZ) induces apoptosis in NTR-mCherry positive epithelial stem cells. (B) The experimental layout for the Controlled Ablation of p63-positive Epithelial Stem Cells (CAPEC) assay. 4dpf zebrafish larvae are treated with MTZ for 4 hours, and then washed out to enable clearance of apoptotic extracellular vesicles, and recovery of the tissue. (C and C’) Treatment with MTZ induces activated-Caspase 3 activation in NTR-positive cells. (D) After MTZ treatment, NTR-positive epithelial stem cells form apoptotic extracellular vesicles (AEVs) over time (hh:mm:ss). (E) The diameter of individual AEVs, with average diameters of 2.413+/− 0.927(SD) µm. (F). An overview of AEV-induced clearance and recovery. E’ depicts an undamaged tissue under homeostatic conditions. Magenta refers to NTR-positive cells, while the green denotes neighboring epithelial stem cells expressing p63:eGFP. F’’ esAEV formation after 4 hours of MTZ treatment. p63-positive cells remain unaffected by MTZ treatment. F’’’ shows the clearance of esAEVs by p63-positive epithelial stem cells. (G) The number of epithelial stem cells engulfing esAEVs at timepoints 8, 12, 18 hours post treatment. N = 15, DMSO 8 and 18 hpt; n = 18, DMSO 12 hpt. n = 32, MTZ 8 and 12 hpt; n = 39, MTZ 18 hpt. ****<0.0001, *0.0140 via a one-way ANOVA with a Tukey’s ad hoc test. (H) Representative images of the compensatory response after MTZ treatment. H’ represents the number of proliferating cells by measuring the amount of BrdU+ cells at 18hpt. n = 38, DMSO. n = 42, MTZ. ****<0.0001 using a two-tailed t-test. Scale bars: C, F, and G = 50 µm, C’ = 10 µm, D = 25 µm.

To identify proteins associated with esAEVs that regulate compensatory proliferation, we purified esAEVs using differential centrifugation and performed proteomic analysis (Figure 2A). Particle size and concentration using tunable resistive pulse sensing identified a fraction enriched in ∼2um AEVs (Figure 2B), consistent with that observed *in vivo* (Figure 1D-E). Liquid chromatography with tandem mass spectroscopy proteomic analysis of the isolated esAEVs identified 421 unique proteins when compared to extracellular vesicles isolated during homeostatic conditions with no apoptosis (Figure 2C). Gene Ontological (GO) analysis defined 14 clusters that had an enrichment score greater than or equal to 1.3, with the highest enrichment scores being pathways involved in cell metabolism (Figure 2D). This analysis also showed that esAEVs are enriched in proteins involved in biological processes such as regeneration, cell-cell junction organization, and lipid modification. We also found several major Damage Associated Molecular Patterns (DAMPs)^36–38^ such as heat-shock proteins (Q90473, and Q645R1), calreticulin (Q6DI13), protein S100 (Q6XG62), and histones H2A and H4 (Q0D272, E7FE07). We also identified proteins such as Angiosinogen (Q502R9), Pro-epidermal growth factor (EGF) (B3DH82), Low-density lipoprotein receptor-related protein (LRP1) (A0A8M2B922) and Galectin-3 (Q6TGN4) which are known to be involved in pathways pertaining to cell proliferation or cell growth (Supplemental Table 1). Finally, we assessed the list for proteins that could play a dual role in stimulating proliferation in epithelial stem cells and regulate inflammatory responses from the immune system. This led to the identification of macrophage migration inhibitory factor (MIF) (F6PCE0). In sum, these data provide new insights into the protein components of esAEVs and their potential role in facilitating compensatory proliferation.

**Figure 2.**
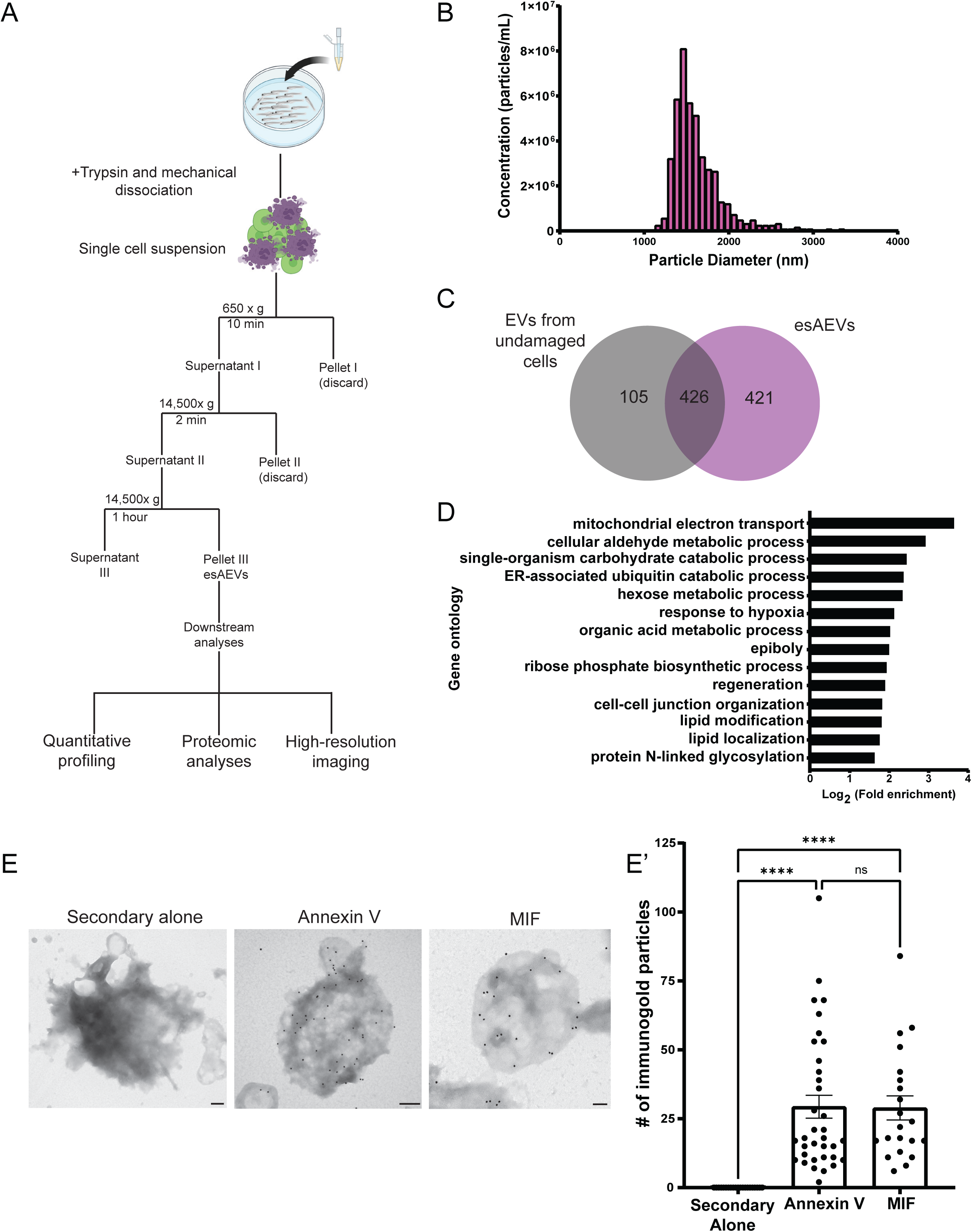
Characterization of purified esAEVs based on protein content and the localization of MIF on the surface. (A) esAEVs are isolated using differential centrifugation. For this study, isolated esAEVs were quantified for size and concentration using qNano technology, submitted for proteomic analysis, and transmission electron microscopy to validate protein localization. (B) A histogram of the size and concentration of isolated esAEVs. Minimum = 1163 nm, Maximum = 3338 nm, Median = 2138 nm, and Mean diameter = 2151 +/− 119.3 (SEM). (C) esAEVs submitted for LC-MS/MS had a total of 925 proteins with 421 being unique to esAEVs compared to the undamaged EV control of 105. (D) Gene ontological analysis of the unique proteins found within esAEVs clustered based on biological process and log fold enrichment. (E) Electron micrographs show positive staining of Annexin V and MIF on the surface of esAEVs. (E’) shows the amount of immunogold particles per AEV. Each point represents the number of immunogold particles per esAEV. The secondary alone measures the number of immunogold particles on 21 esAEVs with a mean particle number of 0.000 +/− 0.000 (SEM). The average number of Annexin immunogold particles is 29.34 +/−4.161 across 35 esAEVs, and MIF has an average of 28.90 +/− 4.373 across 21 esAEVs. The difference between means for Annexin V and MIF is not significant (adjusted p-value = 0.9998). Scale bars: Secondary alone = 200 nm, Annexin V = 200 nm, MIF = 100 nm.

MIF is an attractive target for investigation due to its role in stimulating proliferation, inflammation, and immune cell dynamics. Therefore, we focused our efforts on characterizing the role of MIF in esAEV-mediated signaling. To validate the presence and localization of MIF on purified esAEVs, we assayed for MIF using immunogold labeling transmission electron microscopy. Annexin V served as a control to detect externalized phosphatidylserine (PS) on the surface of esAEVs. We observed no background staining when the gold nanoparticles were administered alone (0.000 +/−0.000). Anti-MIF nanoparticle labeling was applied, it was found to be localized on the surface of esAEVs (28.90 +/− 4.373 nanogold particles) at levels comparable to Annexin V (29.34 +/−4.161 nanogold particles) (Figure 2E). In contrast, the MIF ortholog D-DT (also referred to as MIF-2) displayed little to no detectable nanoparticle localization (Supp. Figure 1). Together, these data suggest that MIF localizes on the surface of esAEVs and could initiate MIF signaling to surrounding epithelial and immune cells.

### esAEVs transporting MIF play a role in regulating epithelial stem cell proliferation

To investigate the role of MIF signaling in apoptosis-induced proliferation and esAEV signaling, we used a combination of pharmacological and genetic approaches to disrupt MIF and its cognate receptor CD74 (Figure 3A). The zebrafish genome contains one copy of *mif*^39^, and two copies of the receptor genes, termed *cd74a* and *cd74b*^40^ (Supp. Figure 2). Larvae expressing NTR in epithelial stem cells (NTR+) were treated with MTZ to induce apoptosis, treated with DMSO (vehicle control), ISO-1^41^, or 4-IPP ^42^ to block MIF signaling, and then incubated with BrdU to label the dividing cells. A compensatory proliferative response was observed with induced apoptosis (MTZ treatment alone compared to DMSO), however we observed a statistically significant decrease in proliferation when induced apoptosis was combined individually with either ISO-1 or 4-IPP treatment to inhibit MIF signaling (Figure 3B-C). To complement the pharmacological approach, we used CRISPR/Cas-9-mediated mutation of *mif*, *cd74a*, and *cd74b*. In the F0 generation of *mif*, *cd74a*, and *cd74b* CRISPR-deleted animals (Supp. Figure 2A-L), or “crispants”^43,44^, we observed a statistically significant decrease in the number of BrdU positive epithelial stem cells (Figure 3D-F). In contrast, F0 larvae injected with guide RNAs for tyrosinase that disrupt pigmentation showed no statistical change in proliferation after induction of apoptosis (Supp. Figure 3A-B). These data support a role for the MIF/CD74 signaling axis in esAEV-induced proliferation.

**Figure 3.**
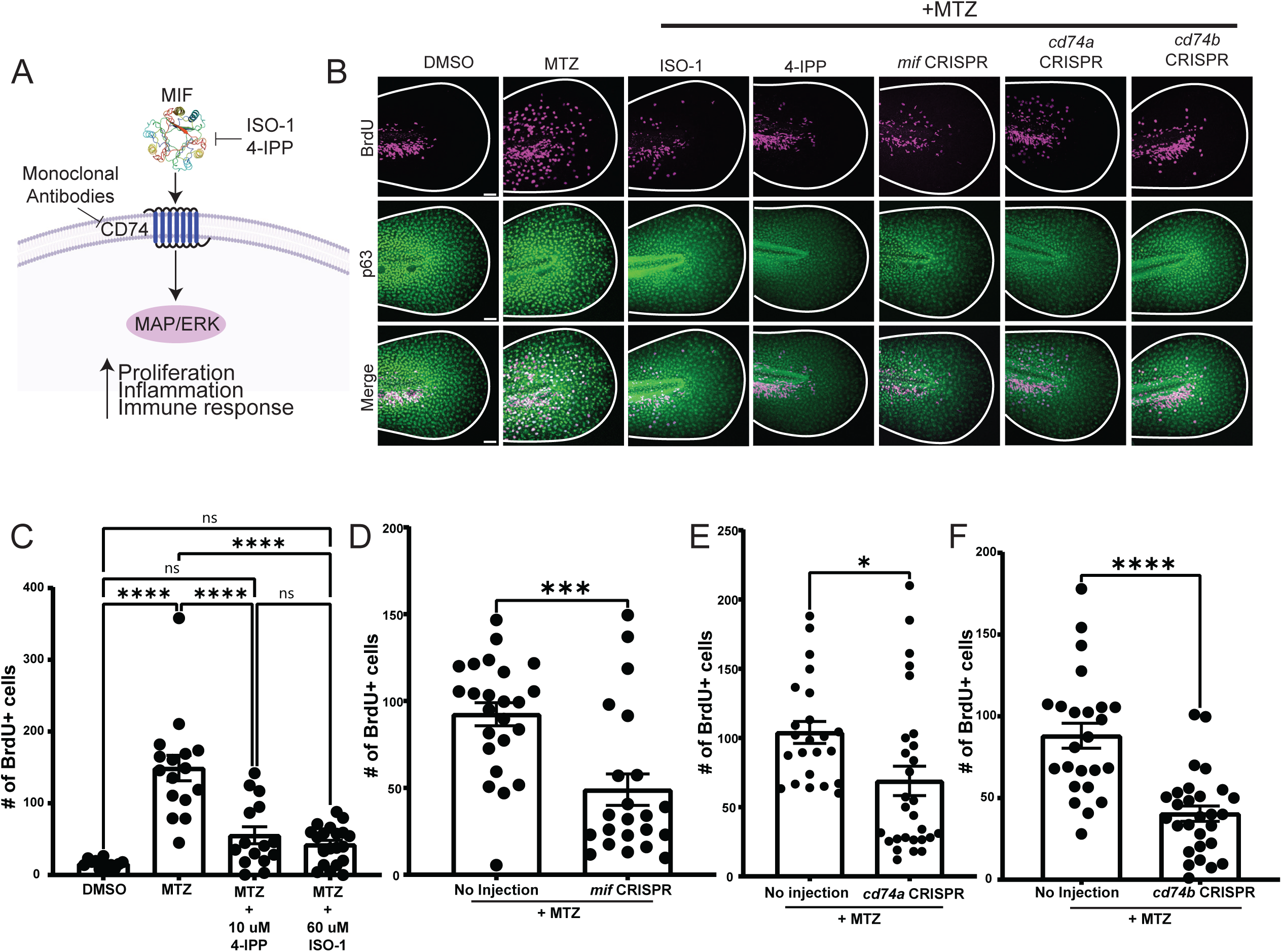
Pharmacological inhibition and genetic manipulation of MIF and its cognate receptor suppresses apoptosis-induced proliferation. (A) An overview of canonical MIF signaling upon binding to its cognate receptor, CD74. (B) Representative images of BrdU staining in p63-positive cells. Size bar = 50 µm. (C) The number of BrdU positive nuclei after MIF inhibition using ISO-1 and 4-IPP before and after apoptosis induction using MTZ. n = 35, DMSO. n = 37 MTZ. n = 40, MTZ + 4-IPP. n = 52, MTZ + ISO −1. **** < 0.0001. (D) Number of BrdU positive nuclei after CRISPR-mediated targeting of *mif*. n= 60 uninjected, n = 62 for *mif* CRISPR across three experiments ***0.0003 using an unpaired two-tailed t-test. (E) Number of BrdU positive nuclei for CRISPR/Cas9 targeting of *cd74a*. n=56 for uninjected, n=54 for *cd74a* CRISPR. *0.0137 using an unpaired t-test. (F) Number of BrdU positive nuclei for *cd74b* CRISPR. n= 62 for no injection, and n= 61 for *cd74b* CRISPR. **** <0.001 using an unpaired two-tailed t-test. All BrdU quantifications were preformed 18 hours post MTZ treatment. These data are presented as means +/− SEM. Scale bar = 50 µm

Our data supports the idea that MIF on the surface of AEVs plays a role in stimulating proliferation to replace cells lost by apoptosis. In addition to its presence on other EV populations^45,46^, MIF has been shown to be either cytosolic or secreted^47^. To test if secreted MIF is contributing to apoptosis-induced proliferation, we first overexpressed human MIF fluorescently tagged with GFP and did not observe a change in epithelial stem cell proliferation (Supp. Figure 5E). Next, we treated larvae with Brefeldin A to disrupt proteins that are secreted through the E.R and found no significant changes in esAEV formation or compensatory proliferation (Supp. Figure 6A-B). MIF can also be secreted via a non-classical secretion pathway that does not involve targeting to the ER^48,49^, and is transported via ABC transporters^50^. Therefore, we also treated larvae with glyburide, a compound that inhibits MIF secretion in THC-1 cells by targeting ABCA1 transport^51^. Glyburide is well tolerated in larval zebrafish without causing alterations to development^52^. After treatment with 25uM glyburide combined with MTZ, we found no significant difference in the number of esAEVs produced or number of BrdU positive epithelial stem cells (Supp. Figure 6C-D). Taken together, these data suggest that MIF delivery on AEVs, rather than by secretion, contributes to compensatory proliferation by the basal epithelial stem cells.

**Figure 4.**
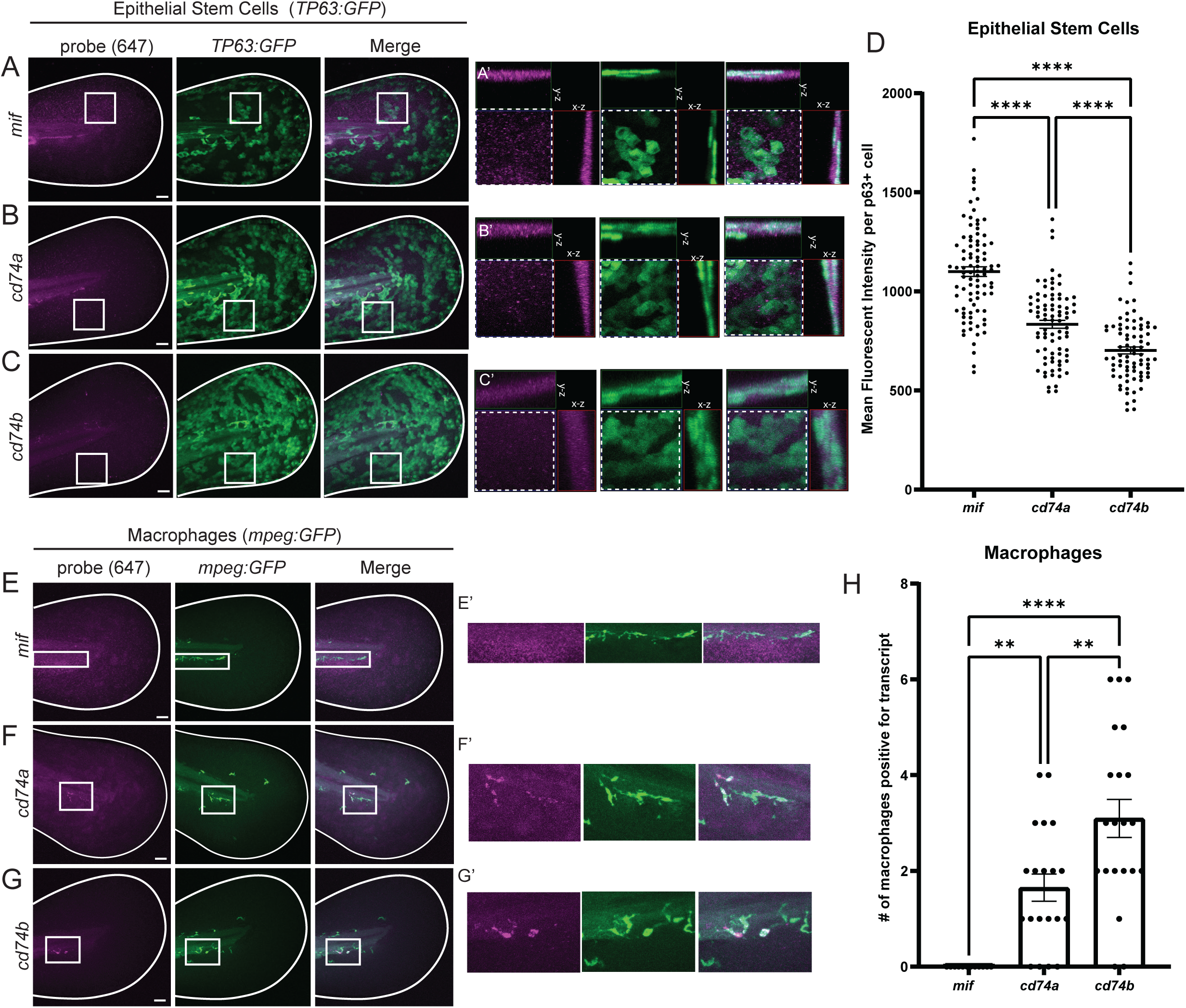
Cell-type specific expression of *mif, cd74a*, and *cd74b*. (A, B, and C) Representative maximum intensity projections of *mif, cd74a,* and *cd74b* fluorescent in situ hybridization probes in the 4dpf larvae expressing GFP in epithelial stem cells (tp63:eGFP). (A’ – C’) are ROI orthogonals showing the presence of *mif, cd74a,* and *cd74b* probe staining the same plane as epithelial stem cells. All probes are imaged in the far-red channel (647nm). (D) Shows the mean fluorescent intensity of *mif, cd74a,* and *cd74b* puncta in individual epithelial stem cells. n = 82, *mif*. n = 89, *cd74a*. n = 79, *cd74b*. **** <0.0001 measured using a One-way ANOVA with a Tukey’s post hoc test. (E, F, G) Demonstrates fluorescent in situ hybridization probes for *mif, cd74a*, and *cd74b* in 4dpf larvae expressing eGFP in macrophages *(mpeg:eGFP)*. E’ shows negative localization of *mif* in macrophages. F’ and G’ shows positive staining of *cd74a* and *cd74b* in macrophages. (H) Shows the number of macrophages per larvae that are *cd74a+, cd74b+,* and *mif+.* n = 20, *cd74a*+. n = 21, *cd74b*+. n = 12, *mif*+. **0.0046 *cd74a* vs. *cd74b*. **0.0057 *cd74a* vs. *mif*. **** <0.0001 *cd74b* vs *mif.* Data are represented as mean +/− SEM. Scale bars = 50 µm.

**Figure 5.**
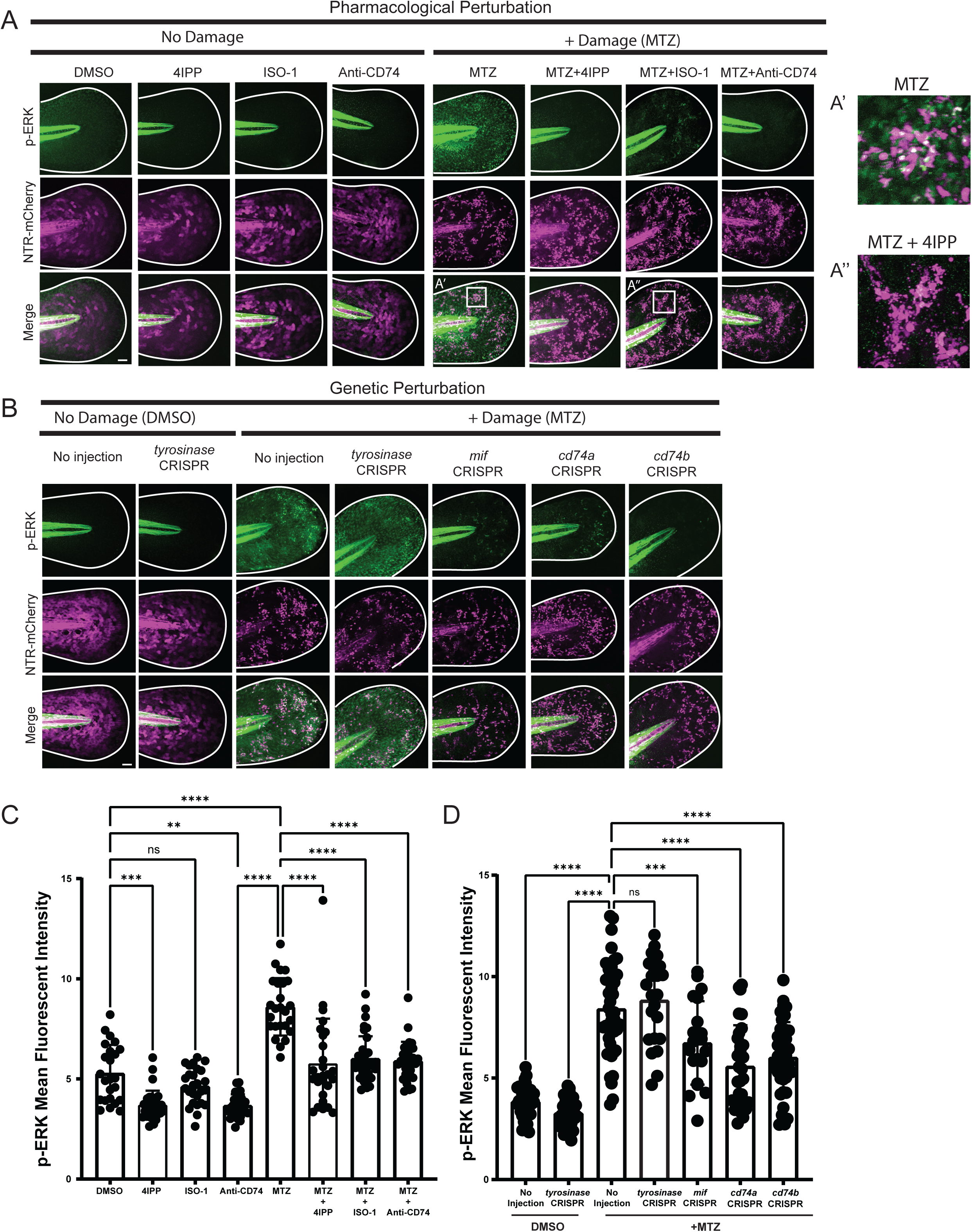
Suppression of MIF and CD74 downregulates p-ERK signaling after apoptosis induction. (A) Representative images of p-ERK signaling with and without damage induction using pharmacological perturbation of MIF/CD74 signaling. A’ Depicts a close up of p-ERK staining within the epithelium after MTZ treatment. A’’ depicts p-ERK staining in the presence of MTZ and 4IPP. (B) Representative images of p-ERK fluorescent intensity after CRISPR targeting of No injection, *tyrosinase (tyr), mif, cd74a,* and *cd74b*. (C) Graph depicting the mean fluorescent intensity of p-ERK across conditions of damage and no damage. DMSO, n=69 ROIs. MTZ, n= 69. MTZ+4IPP, n = 77. 4IPP, n = 71. MTZ+ISO-1, n = 80, ISO-1, n = 73. Anti-CD74, n = 55, MTZ+Anti-CD74, n= 77. For each larvae, 2-3 ROIs within the tail epithelium were selected to measure mean fluorescent intensity. (D) A graph showing the mean fluorescent intensity across CRISPR injected larvae. n = 70, no inj (DMSO). n = 53, *tyr* CRISPR (DMSO). n = 69 no inj (MTZ). n = 44, *tyr* CRISPR (MTZ). n = 40, *mif* CRISPR (MTZ). n = 39, *cd74a* CRISPR (MTZ). n = 40, *cd74b* CRISPR (MTZ). 2-3 ROIs were measured per larvae. Scale bars = 50 µm.

**Figure 6.**
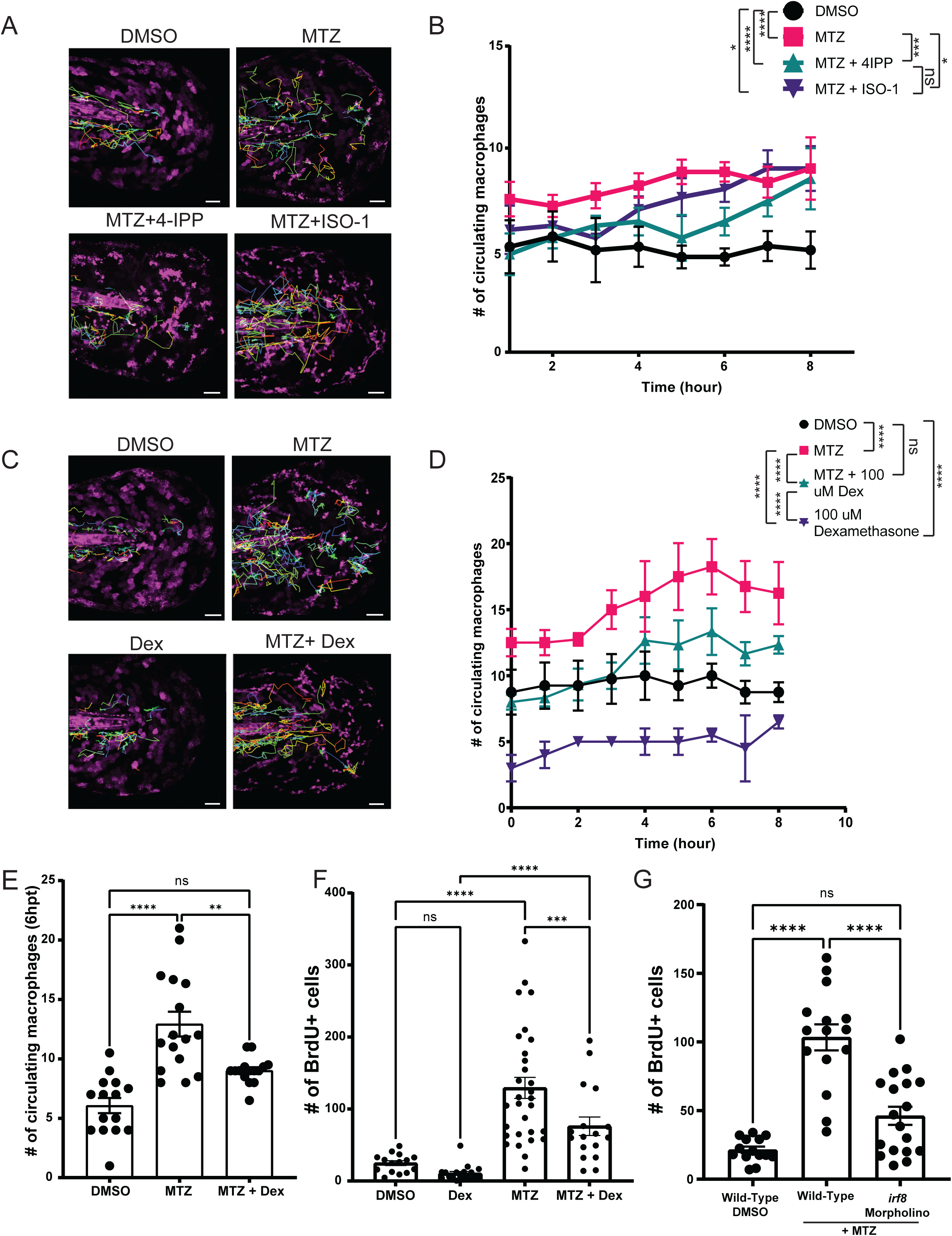
Macrophages play a role in esAEV-induced proliferation. (A) Movement tracking of macrophages over an 8-hour timespan across four treatments denoted by the tracks. (B) Hourly quantification of macrophages extravasating out into the epithelium from the caudal vein based on pharmacological inhibition of MIF. Quantifications are averaged across multiple animals over an 8 hour time period. n = 6 for DMSO and MTZ. n=5 for MTZ+ISO-1, and MTZ+4IPP. * 0.0494, *** 0.0001, **** <0.0001. (C) Movement tracking of macrophages after treatment with dexamethasone to suppress macrophage activity over the course of 8 hours. (D) The number of macrophages surveying areas of apoptotic cells. n=4 DMSO, n=4 MTZ, n=3 for MTZ and Dexamethasone, and n=3 for dexamethasone alone. **** <0.0001, n.s. = 0.1098. (E) Fixed quantifications of macrophage presence at 6hpt with the treatment of dexamethasone. n=19 for DMSO, n = 33 for MTZ, n = 33 for MTZ + 100 uM Dexamethasone. ** 0.0072, **** <0.0001. (F) Assessment of proliferation after treatment of dexamethasone in the presence of esAEVs at 18 hpt. n=41 for DMSO, n=54 for MTZ, n=45 for MTZ + Dexamethasone, and n=43 for Dexamethasone. *** 0.0001, **** <0.0001. (G) The number of proliferating cells after macrophage ablation in the presence of MTZ. n=27 for DMSO, n=41 for Wild-Type + MTZ, and n=45 for IRF8 morpholino + MTZ. **** <0.0001. A two-way ANOVA with a Tukey’s ad hoc test was performed to assess significance for B, D - G. Scale bars = 50 µm.

### esAEVs carrying MIF activate ERK signaling in epithelial stem cells

We next sought to test if the basal epithelial stem cells express both MIF and the CD74 receptors. Hybridization chain reaction (HCR) fluorescent *in situ* hybridization^53^ for *mif* in *TP63:GFP* transgenic animals showed high levels of expression throughout the basal epithelial cells (Figure 4A). In contrast, we observed *cd74a* and *cd74b* were expressed at lower levels within the basal epithelial cells than *mif*, with *cd74a* showing a higher level of expression than *cd74b* (Figure 4B-D). Given that MIF has also been implicated in regulating immune cell function, we also examined expression within macrophages using the *mpeg1:GFP* transgenic line ^54^. Intriguingly, *cd74a* and *cd74b* also appear to be expressed in macrophages (Figure 4E-H). Further, *cd74a* and *cd74b* also localize to macrophages that infiltrate to sites of injury/amputation (Supp. Figure 4A-C). Together, this indicates that esAEVs carrying MIF can signal to CD74a or b on both epithelial stem cells and macrophages.

To determine if MIF signals through CD74 to activate downstream ERK signaling^55^, we analyzed levels of phosphorylated ERK (p-ERK) with and without induced apoptosis, and after disruption of MIF/CD74 signaling. Six hours post esAEV induction, we observed an overall increase in p-ERK fluorescence after induced apoptosis when compared to the DMSO control (Figure 5A,C). Interestingly, p-ERK fluorescence was observed adjacent to the apoptotic cells and esAEVs (Figure 5A’-A’’), suggesting activation in healthy neighboring stem cells. The addition of pharmacological agents ISO-1, 4-IPP, or a humanized blocking antibody against CD74, resulted in a statistically significant decrease in p-ERK fluorescence after induced apoptosis. Further, p-ERK fluorescence was decreased in *mif* and *cd74a* and *cd74b* crispant larvae after induced apoptosis (Figure 5C-D). Conversely, there was no observed increase in p-ERK fluorescence in macrophages at either early (6hr) or later timepoints (10 and 18hr) after induced apoptosis (Supp. Figure 7A-B). Taken together, these data indicate that esAEVs carrying MIF act through interaction with CD74a/b to upregulate p-ERK signaling to stimulate proliferation of epithelial stem cells.

### MIF plays a role in macrophage surveillance activity that contributes to compensatory proliferation in a cell-non-autonomous manner

Inhibition of MIF did not completely decrease apoptosis-induced proliferation, suggesting that other contents of esAEVs play a role in affecting proliferation or there may be another microenvironmental factors contributing to esAEV-induced proliferation. As the name implies, MIF has also been implicated in regulating the migration of macrophages^56,57^. This is in line with our observed expression of *cd74a* and *cd74b* in macrophages (Figure 4E-H). To characterize the response of macrophages after induced apoptosis, we used time-lapse imaging of the *mpeg1:GFP* transgenic line and observed a robust increase in macrophage surveillance activity near the apoptotic cells and AEVs (Figure 6A–B), Supp. Movie 3). An average of 6 engulfment events by the macrophages were observed over 8 hours (Supp. Figure 8A-B), with kinetics that could not support the complete clearance of AEVs from the tissue in this timeframe. These data suggest that macrophages play additional roles beyond solely the clearance of apoptotic corpses and AEVs. Treatment with either 4-IPP or ISO-1 results in decreased macrophage patrolling activity post-induction of apoptosis and AEV formation, with 4-IPP having a stronger suppressive effect than ISO-1 (Figure 6B). The observed decrease in proliferation after ISO/4IPP treatment suggest that the macrophages may also contribute to epithelial stem cell proliferation in a cell non-autonomous manner.

To test if the presence of the immune system contributed to apoptosis-induced proliferation, we treated larvae with the anti-inflammatory dexamethasone to suppress immune cell infiltration after induced apoptosis^58^. Dexamethasone suppressed the number of patrolling macrophages associated with epithelial stem cell apoptosis (Figure 6C-E). Suppression of macrophage infiltration using dexamethasone was also accompanied by significantly decreased compensatory proliferation of TP63+ epithelial stem cells (Figure 6F). To further test this idea, we used antisense morpholino oligonucleotides (MO) to target IRF8, the transcription factor that initiates the myeloid lineage and macrophage formation in early development^59^. *irf8* MO larvae had significantly decreased number of macrophages (Supp. Figure 8C-E), along with reduced epithelial stem cell proliferation after induced apoptosis (Figure 6G). Together, these data support the idea that the presence of macrophages contributes to compensatory epithelial stem cell proliferation after induced apoptosis and AEV formation.

## DISCUSSION

Our studies suggest a role for AEVs derived from epithelial stem cells in stimulating compensatory proliferation after induced apoptosis by modulation of the MIF/CD74 signaling axis. These data support a growing body of research implicating AEVs as key mediators of cell-to-cell communication. For instance, apoptotic bodies have been hypothesized to transfer Wnt3 to stimulate proliferation in neighboring cells during compensatory proliferation in Hydra^7^. Our previous findings using zebrafish has established a role for AEVs carrying Wnt8a to stimulate proliferation in epithelial stem cells^9^. Further, AEVs derived from mouse mesenchymal stem cells stimulate proliferation of mesenchymal stem cells through transfer of molecules that can upregulate the Wnt/β-catenin signaling pathway^8^. Additional effects of AEVs have been observed in stem cell populations such as endothelial progenitor cell differentiation^60^, mononuclear osteoclast progenitors^61^, and cardiac precursor cells^62^. Overall, these studies suggest that AEVs play an important role in regulating stem cell proliferation and tissue regeneration. Our current studies extends these findings further by providing an *in vivo* assessment of AEV activity in conjunction with an intact innate immune system.

A key question from these studies is how putative signals are transferred from apoptotic cells to neighboring healthy cells to initiate compensatory proliferation. Our studies suggest a critical role for surface-localized MIF on AEVs in mediating intercellular communication from dying epithelial stem cells to healthy neighboring epithelial stem cells and macrophages. Extracellular vesicles, including both various EV subtypes and exosomes, can transfer DNA, mRNA, and proteins to facilitate short-range communication between cells^63^. Mesenchymal stem cell exosomes carrying MIF have also been shown to enhance myocardial repair, promote angiogenesis and reduce fibrosis in the heart^64^. Similarly, exosomal MIF from nasopharyngeal carcinoma promotes metastasis by enhancing macrophage survival^46^. These findings suggest that AEVs carrying MIF also play important roles in regulating diverse biological processes, including immune responses, tissue repair, and cancer progression.

The interaction between the cytokine MIF and its receptor CD74 can trigger downstream signaling cascades such as ERK1/2 to drive a change in cellular behaviors including proliferation, migration, and survival. In our study, we showed that AEVs promote an increase in p-ERK signaling in neighboring epithelial stem cells to drive proliferation. MIF has also been shown to drive proliferation and migration of airway muscle cells^65^, spermatogonial cells^66^, dendritic cells^67^ in an ERK1/2 dependent fashion. While we did not observe a change in p-ERK in macrophages, other studies have shown that EVs carrying MIF can stimulate p-ERK in macrophages^68^, promoting their activation and modulating their immune responses. A possible explanation for this is that CD74 can recruit additional co-receptors, such as CXCR2^29^ or CXCR4^69^, instead of CD44^70^ in a MIF dependent manner and activate other downstream signaling events such as PI3K/Akt, and calcium dependent integrin activity^29^. What mediates the particular co-receptors that CD74 recruits is not well understood, and future characterization of these interactions may inform which downstream pathways are taking place in macrophages to facilitate compensatory proliferation.

Apoptosis has traditionally been regarded as an immunologically silent form of cell death that resolves with minimal induction of inflammation^71^. The rapid externalization of PS during apoptosis serves as an ‘eat-me’ signal and exhibits anti-inflammatory properties^72^. DAMPs are released during apoptosis and cellular stress events^73^, calling this previously held dogma into question^38^. Intriguingly, AEVs can also transport DAMPs such as HMGB1^20^, a molecule involved in tissue repair^74^. Other extracellular vesicle populations such as small EVs and microvesicles can carry a variety of DAMPs^36^, with HSP70 being localized on the surface of exosomes^75^. MIF can be secreted by a variety of cell types in response to tissue damage, infection, and other forms of stress, and has been shown to have pro-inflammatory and immune-regulatory effects^47,76^. We did not observe an impact on proliferation with pharmacological inhibition of MIF secretion, or with genetic overexpression of human MIF, suggesting that apoptosis is critical for the ability of MIF to promote a compensatory proliferation response. Therefore, MIF may represent a novel DAMP with unique properties and functions.

Our studies indicate that macrophages contribute to AEV-mediated compensatory proliferation. We observed expression of *cd74a* and *cd74b* in macrophages, and established a role for AEV delivered MIF in promoting macrophage surveillance activity after induction of apoptosis. Importantly, dampening of inflammation via treatment with dexamethasone or depletion of the macrophage population decreased proliferation, suggesting a cell-non autonomous mechanism for the stimulation of proliferation during the re-establishment of epithelial tissue homeostasis. This is echoed by studies showing that macrophages play a key role in facilitating tissue repair^77^, and that attenuation of macrophages impairs wound healing and tissue regeneration^78–81^. Apoptotic cells can prime macrophages toward a more “pro-regenerative” M2 phenotype^82^, and M2 macrophages are known to secrete a variety of signals following injury^78^. Recent work has highlighted that MIF can induce macrophages to secrete mmp-9 downstream of ERK1/2 signaling^83^. The types of signals macrophages secrete in response to AEV’s carrying MIF remains unclear. Identification of the signals produced by macrophages will be an interesting topic for future studies of compensatory proliferation.

In summary, our studies define a role for MIF carried by AEVs in the reestablishment of tissue homeostasis in a dynamic process that engages both epithelial stem cells and macrophages. We propose that AEVs carrying MIF play a dual role in sustaining homeostatic cell numbers, to directly stimulate epithelial stem cell repopulation and guide macrophage behavior to cell non-autonomously contribute to localized proliferation.

## MATERIALS AND METHODS

### Zebrafish handling and husbandry

Adult Zebrafish were maintained at the MD Anderson Cancer Center fish facility in accordance with the institutional guidelines and best practices for animal care. Zebrafish embryos were maintained at 28°C in E3 medium.

### Transgenic lines

The Basal-GET line was used to drive the expression of nitroreductase in epithelial stem cells: *Et(Gal4-VP16)^zc^*^1036^*^A^, Tg(UAS-1b:nsfB-mCherry)^c^*^264^. The Basal-GET line was crossed with a *Tg(mpeg1:GFP)* line^54^ to visualize macrophage dynamics in the presence of apoptotic cells. The *Tg(tp63:GFP)*^21^ line was used to visualize epithelial stem cells.

All transgenic lines were maintained on a wild-type AB background.

### Controlled Ablation of p63-positive epithelial stem cells (CAPEC) assay

To temporally and spatially ablate p63-positive epithelial stem cells, we used the controlled ablation of p63-positive epithelial stem cells (CAPEC) assay. The Basal-GET line, which expresses nitroreductase-mCherry in basal cells, was treated with 10 mM metronidazole (MTZ) for four hours. This treatment induced the production of epithelial stem cell-derived extracellular vesicles (esAEVs) in NTR-mCherry-positive cells, which were further studied through live imaging or post-recovery analysis.

To assess proliferation or recovery phenotypes, larvae were subjected to an 18-hour recovery time prior to bromodeoxyuridine (BrdU) incorporation. After the recovery period, BrdU was administered to label dividing cells, allowing us to analyze proliferation and recovery phenotypes in response to stem cell ablation.

### Isolation of esAEVs and quantitative analysis

After inducing apoptosis in epithelial stem cells in 4dpf larvae, a combination of mechanical dissociation, trypsonization, and centrifugation were used to isolate esAEVs. After a 4-5 hour treatment with MTZ, 150-200 Basal-GET NTR+ larvae were transferred to a 1.5mL eppendorf tube, and washed once with tissue-culture grade 1X PBS. Once the larvae settled in the tube, 1mL of pre-warmed 37°C Trypsin was added to the tube. A scalpel was used to chop the larvae for roughly 60-90 seconds. The larvae were then placed on a nutator to rock gently for 10 minutes. Chopping and placement on the nutator was repeated twice more. The larvae were then placed in a 4°C centrifuge for 10 minutes at 650 x g. The supernatant was transferred to a new tube and centrifuged for 2 minutes at 14,500 x g, 4°C to pellet large cells. The supernatant was transferred to a new tube and centrifuged for 1 hour at 14,500 g, 4°C. In this study, tunable-resisitive pulse sensing (qNano, Izon) was used to quantify the size and concentration of esAEVs. For the qNano, CPC 2000 calibration particles were used with an NP 2000 pore. esAEVs were diluted 1:50 in measurement electrolyte prior to measuring the size and concentration. Look to Figure 2A for a visual representation of this methodology.

### Proteomic profiling of esAEVs

Proteins were isolated from esAEVs and run on a gel. Coomassie gel pieces were washed, de-stained and digested in-gel with 200 ng modified trypsin (sequencing grade, Promega) and Rapigest (TM, Waters Corp.) for 18 hrs at 37°C. In-solution samples were precipitated with 5:1 v/v of cold acetone at −20°C for 18 hrs, then centrifuged and the acetone was removed prior to treatment with Rapigest (100°C for 10 min) followed by addition of trypsin. Resulting peptides were extracted and analyzed by high-sensitivity LC-MS/MS on an Orbitrap Fusion mass spectrometer (Thermo Scientific, Waltham MA). Proteins were identified by database searching of the fragment spectra against the SwissProt (EBI) protein database using Mascot (v 2.6, Matrix Science, London, UK) and Proteome Discoverer (v 2.2, Thermo Scientific). Peptides were subject to 1% FDR using reverse-database searching.

### Gene ontology analysis of proteins unique to esAEVs

GO enrichment analysis was performed using DAVID^84^. The 421 unique proteins to esAEVs were uploaded to DAVID, where the following criteria was applied: Using the highest stringency for biological processes, a total of 72 clusters with different enrichment scores were returned. Applying an exclusion criterion of no less than 1.3 for enrichment scores, 14 clusters were appropriate for analysis. Within each cluster, terms with fold enrichments greater than 1.5 and the lowest false discovery rate (FDR) were further selected as biologically relevant.

### Transmission electron microscopy and Immunogold labeling

With the assistance of the M.D. Anderson Cancer Center Electron Microscopy Core Facility, isolated AEVs were submitted for immunogold labeling. AEVs were fixed in 2% Glutaraldehyde in 0.1M PBS, pH 7.5. In order to visualize the membrane, esAEVs were whole-mounted. During the whole mount, AEVs were mildly permeabilized with saponin to expose surface antigens. Primary antibodies were administered at the following concentrations: Annexin V (1:200), MIF (1:100), and D-DT (1:200). The secondary antibody was administered as a 1:20 dilution of gold nanoparticles.

### Pharmacological treatments

All pharmacological treatments were performed for four hours unless otherwise specified. All drugs were washed out 3 times with E3. Metronidazole (MTZ) (Sigma, M3761) was made fresh daily as a 1M stock in DMSO stored at room temperature and is protected from light. MTZ was added to E3 medium at a concentration of 10mM. ISO-1 (Tocris, 4288), 4-IPP (Tocris, 3429), and Brefeldin A (Millipore-Sigma, B7651-5MG) were dissolved in DMSO and stored at 10mM stocks at −20°C. ISO-1 was added to E3 medium at a concentration of 60uM, 4-IPP and Brefeldin A are added at a concentration of 10uM. Dexamethasone (Tocris, 1126) was stored at −20°C as a 100mM stock, and replaced on a monthly basis. 4dpf larvae were treated at a concentration of 100uM. The Human Anti-CD74 antibody (BioLegend, 326802) was stored at 4°C, and added to E3 at a 1:400 dilution.

### Microscopy

A Zeiss LSM800 Laser Scanning confocal microscope was used for movie and image acquisition. Images were acquired as z-stacks using 20x for tail tips. 10x objective was used to acquire tiled region images of entire larvae and stitched together to present a full image. A Zeiss Axiozoom fluorescent microscope was used for acquisition of images for the BrdU immunohistochemical analyses. All microscopy images were processed using Zen software. Any adjustments made to brightness and contrast were applied consistently across images. The resulting images were then used for further analysis and quantification.

### Larvae fixation and fluorescent immunostaining

After treatment or recovery, larvae were fixed in 4% formaldehyde (Sigma) in 0.05% PBST. Larvae were left on gentle rocking overnight at 4 degrees. The fixative was washed out using 6 x 5 minute washes in PBST 0.5%. BrdU detection was done by adding 2N Hydrochloric acid in ddH2O for 45 minutes under gentle rocking at room temperature. The HCL was washed out with 6 x 5 minute washes in PBST 0.5%. Blocking was performed using 10% goat serum in blocking buffer for 1-2 hours. Antibodies were diluted in blocking solution, and used to stain larvae overnight in 4°C. The primary antibody was washed out with 6 x 20 minute PBST 0.5% washes. Larvae were blocked 1-2 hours before the secondary antibody was added. Larvae were left overnight in the dark at 4 degrees. The secondary was washed out using 6 x 20 minute PBST 0.5% washes. Body parts just above the cloaca were removed and discarded during mounting. 80% glycerol in PBS was used to preserve the fluorescence of larvae.

### Antibodies

1:200 Rat anti-Brdu (AbCam, ab6326), 1:200 Rabbit anti-Activated Caspase 3 (BD Biosciences, 559565), 1:200 Mouse anti-Phospho-p44/41 (ERK1/2) (Cell Signaling, Antibody #9101), 1:1000 DAPI, 1:200 Rabbit anti-eGFP antibody (ThermoFisher Scientific, OSE00002W), Rabbit anti-Annexin V ( AbCam, ab14196), Rabbit anti-MIF (AbCam, ab65869), Rabbit anti-DDT (AbCam, ab115785).

### BrdU incorporation and detection

BrdU incorporation was performed using 10mM BrdU, and 5% DMSO in E3 at 18 hpt for 45 minutes. After incorporation, animals were washed 3x with E3, left to recover for 45 minutes in E3 alone, and then fixed in paraformaldehyde overnight at 4°C. We detected BrdU positive cells after cell permeabilization using 2N HCL for 45 minutes. The staining protocol proceeded with applying monoclonal rat anti-BrdU primary antibody followed by the appropriate secondary antibody.

### Immunostaining and Quantification of p-ERK signal

4dpf Basal-GET larvae were drug treated with MTZ, and different agents to perturb MIF activity for 4 hours. Larvae were fixed 6 hours post treatment, and stained for p-ERK (1:200 dilution) (Cell Signaling, Antibody #9101). All p-ERK signal was visualized in the far-red channel as this has the least autofluorescence in the presence of apoptotic cells. The tail epithelium of the larvae were imaged with the exact same imaging parameters. 100um x 100 um regions of interest were drawn around NTR-mCherry cells in control and damage conditions. Tails had a maximum of three ROIs. The mean fluorescent intensity was recorded across a z-stack within each ROI.

### Fluorescent *in situ* hybridization

Probes for the *in situ* hybridization were purchased from Molecular Instruments, inc. The genes and the accession numbers provided to Molecular Instruments are as follows:

MIF (NM_001043321.1), CD74a (NM_131590.1), CD74b (NM_131372.2).

MIF was detected using a B1 amplifier while CD74a and CD74b were detected with a B3 amplifier. Larvae were stained according to manufacturer detection and staining procedures^53^, with modifications outlined by Ruiz et al.^85^ All larvae were imaged in the exact same conditions to allow for comparisons between probes, and all imaging was performed using the far-red channel to minimize background fluorescence.

### Generation of CRISPR-edited larvae

The program CHOP-CHOP^86,87^ was used to design highly specific guides directed against MIF, CD74a, and CD74b. 1-cell stage embryos were injected with two guides for each gene with Cas9 protein (NEB, M0646T). Tyrosinase was an injection control to validate the efficacy of the Cas9 protein. Embryos were raised to 4dpf for further experimentation. We validated the efficiency and accuracy of Cas9 by using Sanger sequencing, followed by TIDE^88^ and ICE^89^ (Synthego Corporation) analysis to compare the signal traces between edited and unedited animals at the expected cut sites.

### gRNA sequences

*tyrosinase*: 5’-GGACTGGAGGACTTCTGGGG**AGG**-3’

*mif* gRNA 1: 5’-TGAGCGAGCAGAGCGCACAC**GGG**-3’

*mif* gRNA 2: 5’-TGCTAAAGACTCGGTTCCGG**CGG**-3’

*cd74a* gRNA 1: 5’-TCCTGGGTCGAGGTGATGCA**AGG**-3’

*cd74a* gRNA 2: 5’-GCTGAATCAGAGACTCGTTC**TGG**-3’

*cd74b* gRNA 1: 5’-TTAACATGGGACCTCAGCCA**AGG**-3’

*cd74b* gRNA 2: 5’-GGCGGTCTCCTCGTCTCTCC**AGG**-3’

### Morpholino Oligonucleotides

The morpholinos used in this study were obtained from GeneTools, and the sequences were obtained from ZFIN (https://zfin.org/). The injection scheme for Irf8 was modeled after Madigan et al.^90^ A GeneTools 25 nucleotide scrambled (Random) control oligo was used as an injection and morpholino control.

### Irf8 morpholino: AATGTTTCGCTTACTTTGAAAATGG <u>Generation of *hsp70:MIF-turboGFP* larvae</u>

The Human cDNA construct of MIF-turboGFP was obtained from origene (RG205106). Overlap PCR was used to flank both ends of MIF cDNA with ATTB sites, and the AttB flanked MIF-turboGFP was inserted in pDONR221. The entry vector, 3’ hsp70 element, middle entry clone, and 5’ middle entry clone were combined in equimolar amounts with LR clonase overnight at room temperature to generate a transgenesis vector to be transformed and grown in carbenicillin plates. Transposase vector was linearized using NotI, and in vitro transcribed using mMessage. The *hsp70:MIF-turboGFP* larvae were mixed with transposase to concentrations of 7 ng/uL of vector and 50ng of transposase mRNA into Basal-GET embryos. The hsp70 constructs were gifts from the Rosa Uribe lab. The destination vector, polyA tail, and transposase plasmid were gifts from the Kristen Kwan lab.

### Heat shock induction of MIF induction in larval zebrafish

Larvae were heat-shocked twice at 37°C for one hour at 3dpf approximately 24 hours and 16 hours before treatment at 4dpf. Larvae expressing MIF-GFP and NTR-mcherry underwent MTZ treatment. Proliferation (via BrdU incorporation) was assessed at 5dpf following recovery and prolonged MIF expression.

### Tracking and quantification of macrophage movement

Using a Basal-GET line crossed to *Tg(mpeg:GFP)*, macrophages were imaged every five minutes using a Zeiss LSM 800 confocal microscope. The number of macrophages outside the notochord and present in the epithelial tissue were manually counted every hour, and graphed over time using GraphPad Prism. Imaris was used to track the overall movement of macrophages over an 8-10 hour time period.

### Software

Zeiss Zen Blue was used to analyze the mean fluorescent intensity of p-ERK signal, and fluorescent *in situ* hybridization. Zen blue was also used to quantify the number of BrdU+ nuclei. GraphPad Prism (version 9) was used for statistical analysis and graph generation. DAVID was used for protein Gene Ontology analysis. Imaris (version 9) was used to track the movement of macrophages in the larval zebrafish tail epithelium.

### Statistical analysis

Statistical analysis was performed using GraphPad Prism version 9. A t-test was used to assess significance between two groups, and an ANOVA was used to determine the significance between three or more groups with a Tukey test as the ad hoc analysis. Results are reported with the standard error of the mean unless specified. A p-value less than 0.05 was considered statistically significant.

## ACKNOWLEDGEMENTS

These studies were supported by National Institutes of General Medical Sciences, R01GM124043 and R35GM149226 to GTE. We thank Kenneth Dunner from the M.D. Anderson High Resolution Electron Microscopy (HREM) facility for assistance with transmission electron microscopy and the immunogold labeling. The authors acknowledge the support of the HREM facility by the CCSG grant NIH P30CA016672, and the UT-M.D. Anderson Proteomics Facility by the M.D. Anderson Cancer Center and the NIH High-End Instrumentation program grant 1S10OD012304-01. We would like to thank Drs. Kristen Kwan, Ghislain Breton, and Rosa Uribe for gifting genetic constructs for transgenesis. We sincerely appreciate Philip Kahan for his care of the adult zebrafish that made this study possible.

## AUTHOR CONTRIBUTIONS

S.E. and G.T.E conceived of the methodology and planned the experiments. S.E. and I.A. performed all the experiments and the subsequent analysis. S.E. and G.T.E wrote the manuscript.

## SUPPLEMENTAL FIGURE LEGENDS

**Supplemental Figure 1.**
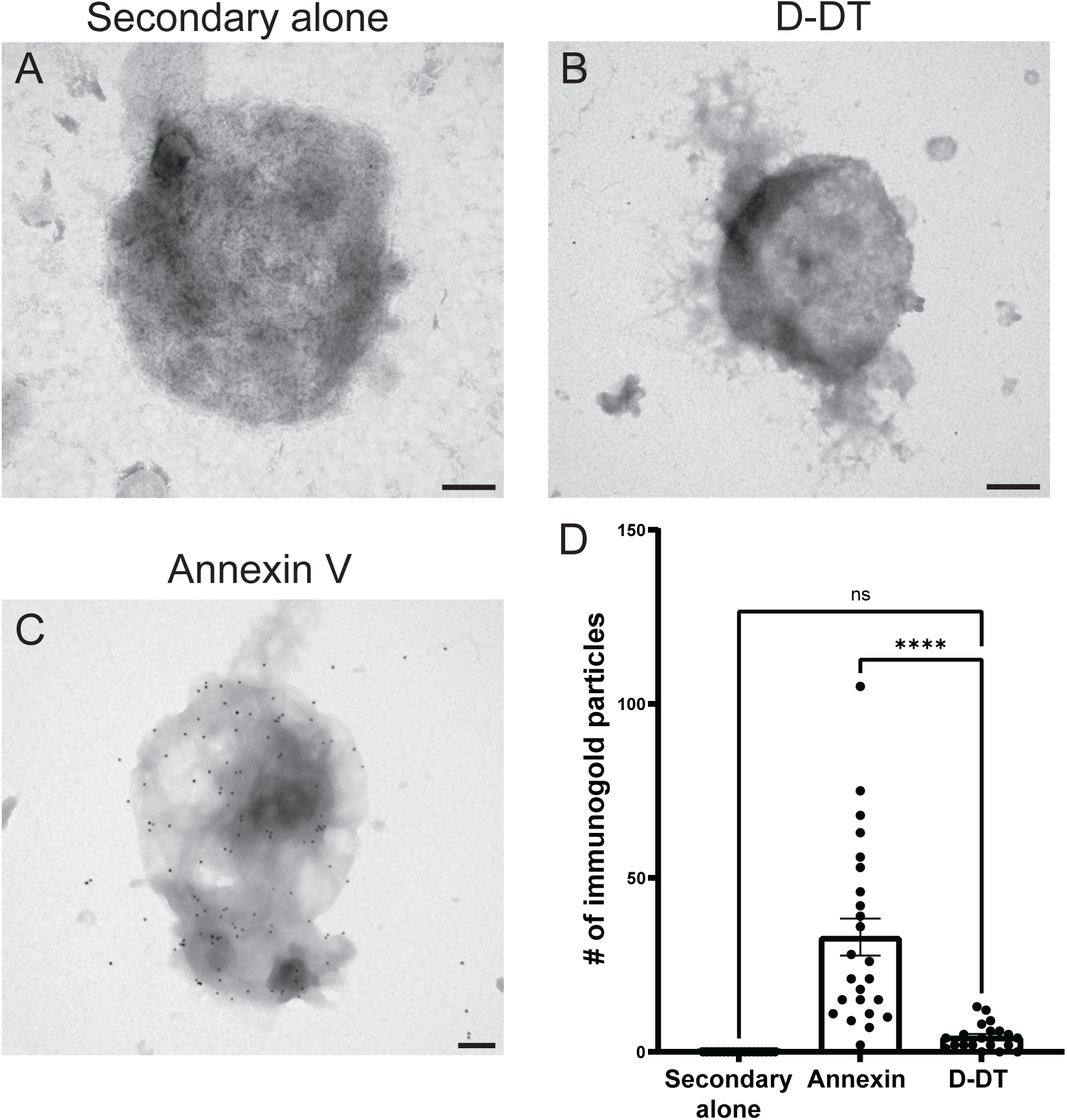
D-DT (MIF-2) is not localized to the surfaces of esAEVs. (A) A representative image of an esAEV administered a control stain of Secondary Alone (also referred to as secondary alone). Size bar = 200 nm. (B) D-DT staining on an esAEV. Scale bar = 400 nm (C) Annexin staining on esAEV. n = 200 nm. (D) Quantitative representation of nanogold particles for Secondary Alone (mean+/−SEM = 0.000 +/− 0.000), D-DT (mean = 4.350 +/− 0.8438), and Annexin V (mean = 33.00 +/− 5.310). Each dot represents an individual esAEV as represented in panels A-C. n = 18, Secondary Alone. n = 24, Annexin V. n = 20, D-DT. The difference between means for Annexin V and D-DT was significant (adjusted p-value<0.0001). There was not a statistically significant difference in means between D-DT and Secondary alone (p = 0.6939).

**Supplemental Figure 2.**
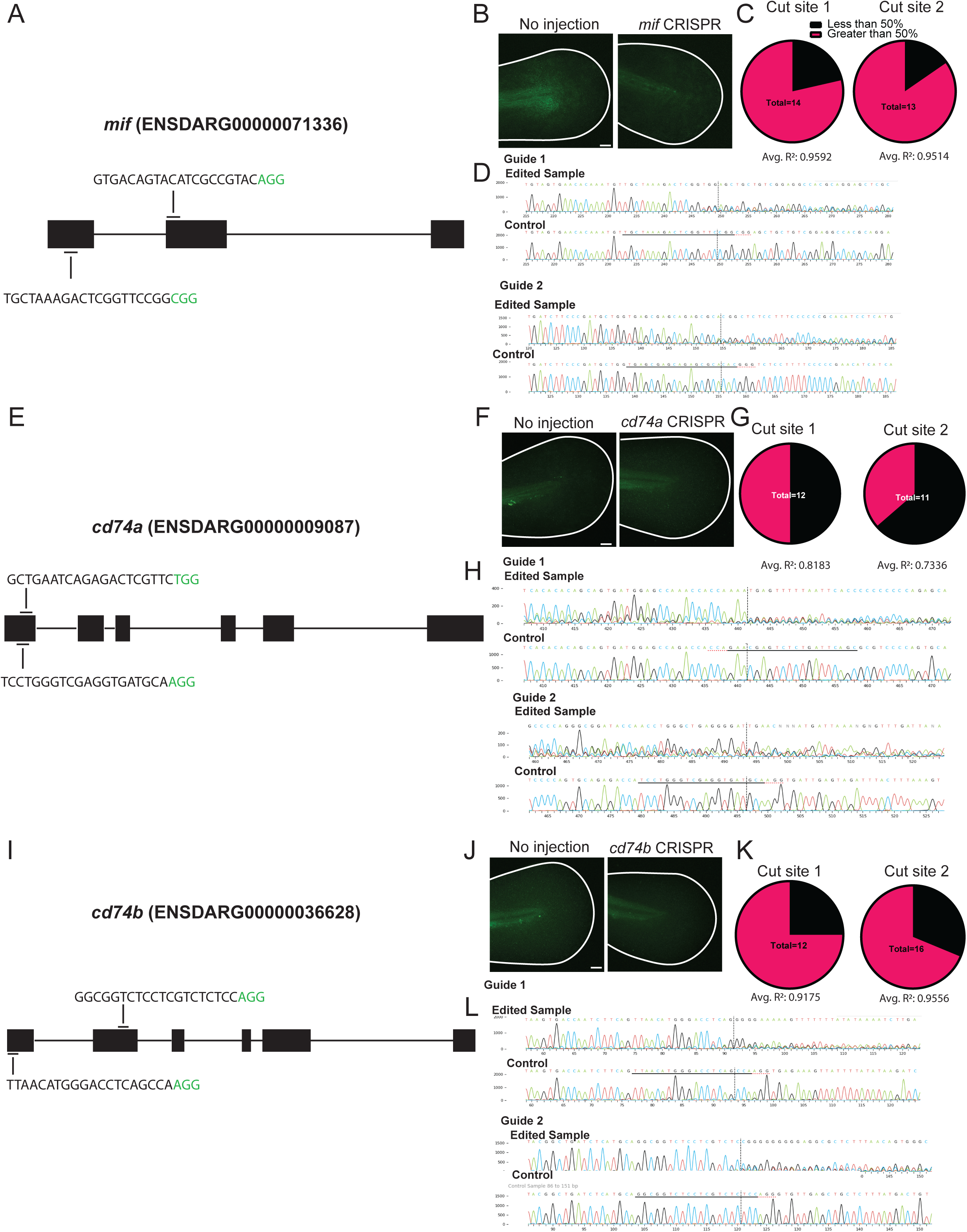
Genotyping of F0 CRISPR-edited larvae using sanger sequencing and fluorescent *in situ* hybridization (FISH). (A, E, I) The gene structures for *mif*, *cd74a*, and *cd74b*. The sgRNAs for each gene are represented along with the general locations within each respective gene. The green text highlights the PAM sequences. The text in the parenthesis are the ensembl genome browser accession numbers, and the information that was entered into CHOP-CHOP to design sgRNAs. (B, F, J) FISH staining of uninjected vs *mif*, *cd74a*, and *cd74b* CRISPR larvae respectively. (C, G, K) The comparison CRISPR editing for two cut sites for *mif*, *cd74a*, and *cd74b*. Magenta represents cutting efficiencies greater than 50%, and black represent less than 50% cutting efficiency as predicted by TIDE or ICE analysis. R-squared values represent the sequence alignment between control and CRISPR-edited samples. (D, H, L) Representative signal traces comparing control to CRISPR edited larvae upstream of the PAM site for *mif* (B), *cd74a* (F), and *cd74b* (J). Scale bars = 50 µm.

**Supplemental Figure 3.**
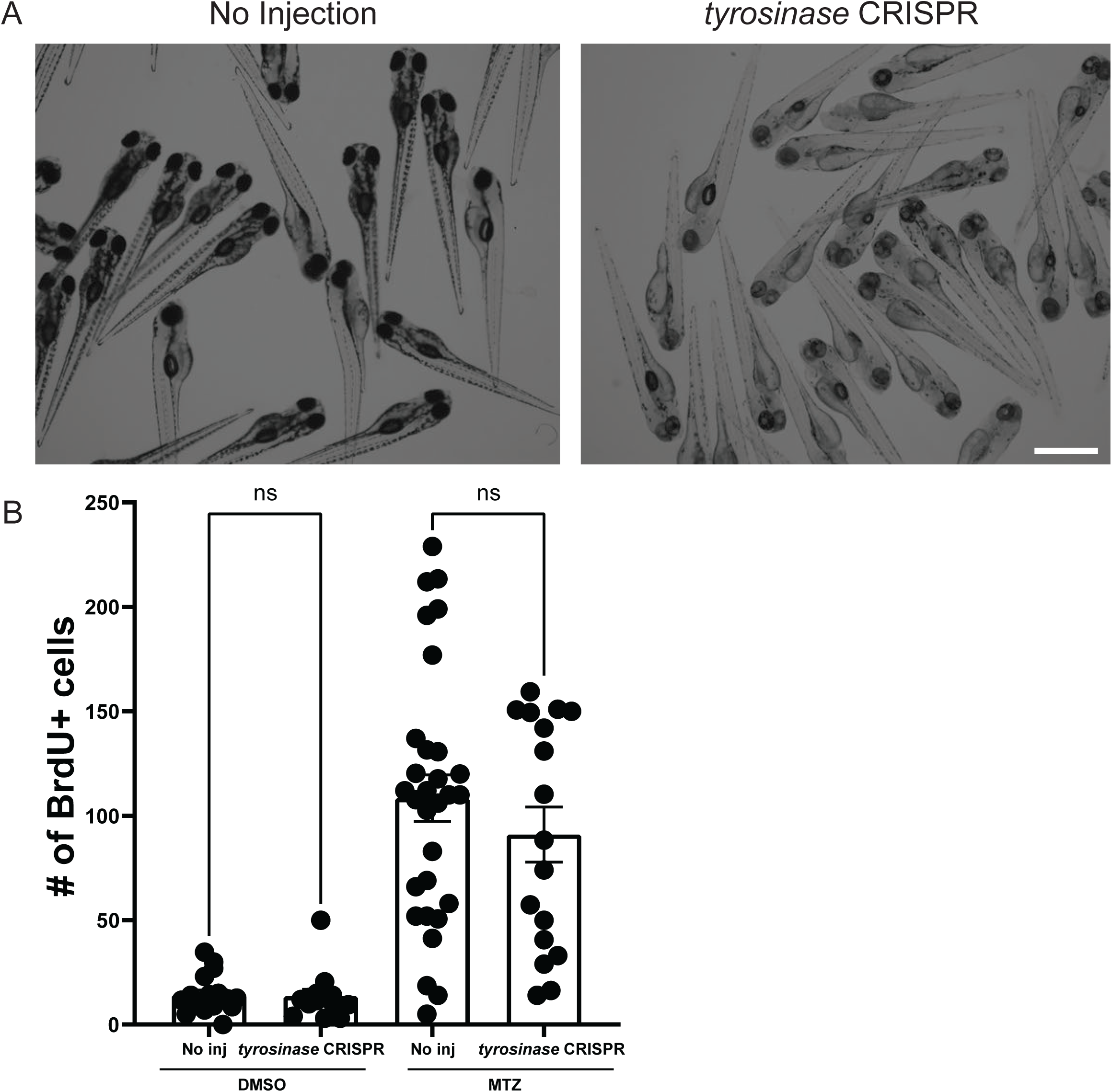
CRISPR/Cas9 editing of *tyrosinase* does not affect apoptosis-induced proliferation. (A) Phenotypic differences between no injection and CRISPR-edited *tyrosinase* larvae. The loss of pigment is a visual confirmation of successful *tyrosinase* gene editing, and this serves as a proxy for the functionality of the Cas9 protein. (B) Assessment of apoptosis-induced proliferation between uninjected larvae and *tyrosinase* CRISPR larvae after the addition of MTZ. A two-way ANOVA demonstrates that there is no statistical difference between the means for MTZ-treated uninjected and *tyrosinase* CRISPR. n = 41, uninjected, DMSO. n = 59 uninjected, MTZ. n = 23 *tyrosinase* CRISPR, DMSO. n = 40 *tyrosinase* CRISPR, MTZ. Scale bar = 1000 µm.

**Supplemental Figure 4.**
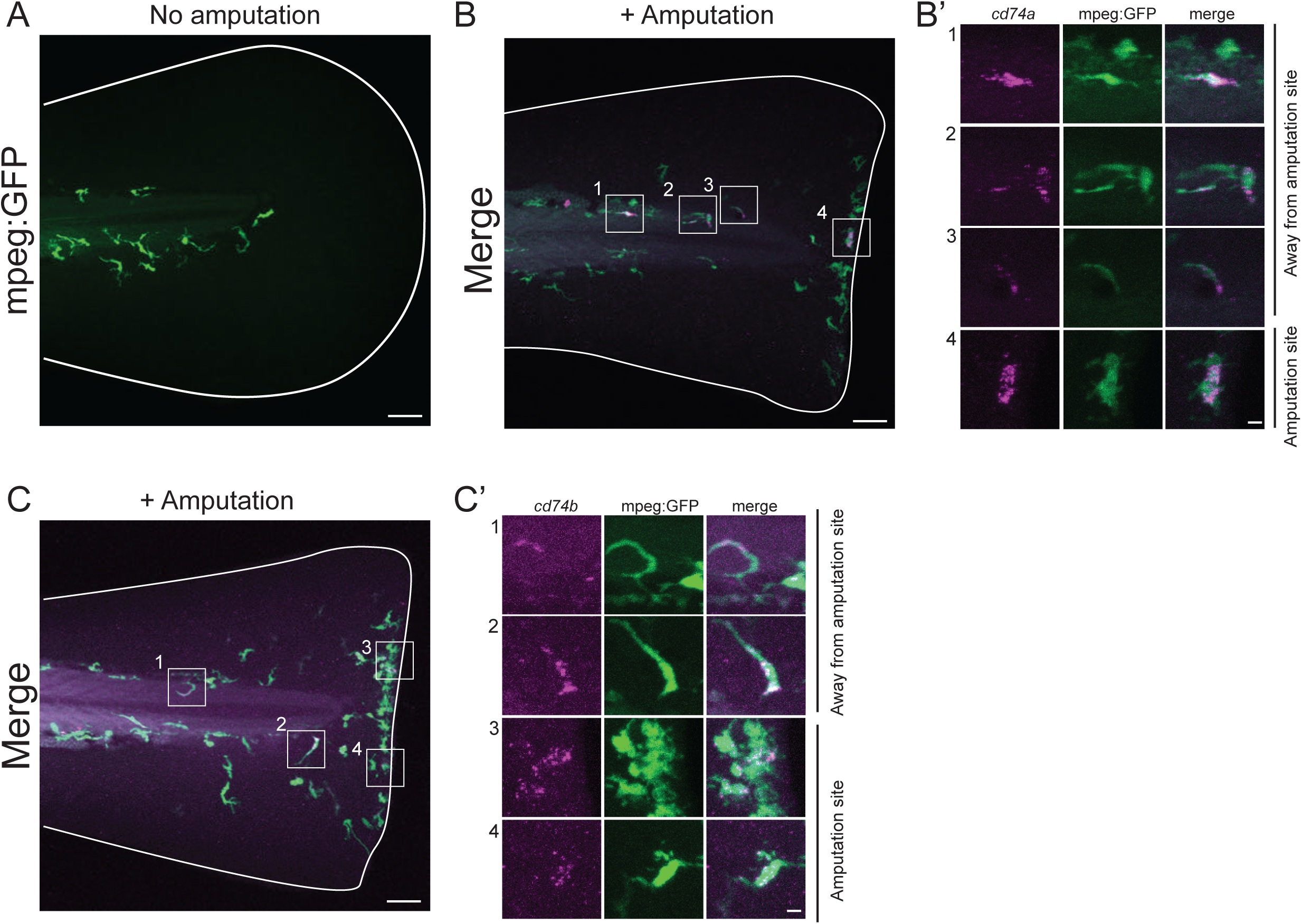
c*d*74a and *cd74b* expression in macrophages. (A) A representative image of macrophage location during homeostatic conditions in an *mpeg:GFP* transgenic line. (B) A representative image of *cd74a* transcripts in macrophages (B’) at and away from the amputation site. (C) A representative image of *cd74b* transcripts in macrophages (C’) at and away from the amputation site. Scale bars: A, B, and C = 50 µm, B’ and C’ = 5 µm.

**Supplemental Figure 5.**
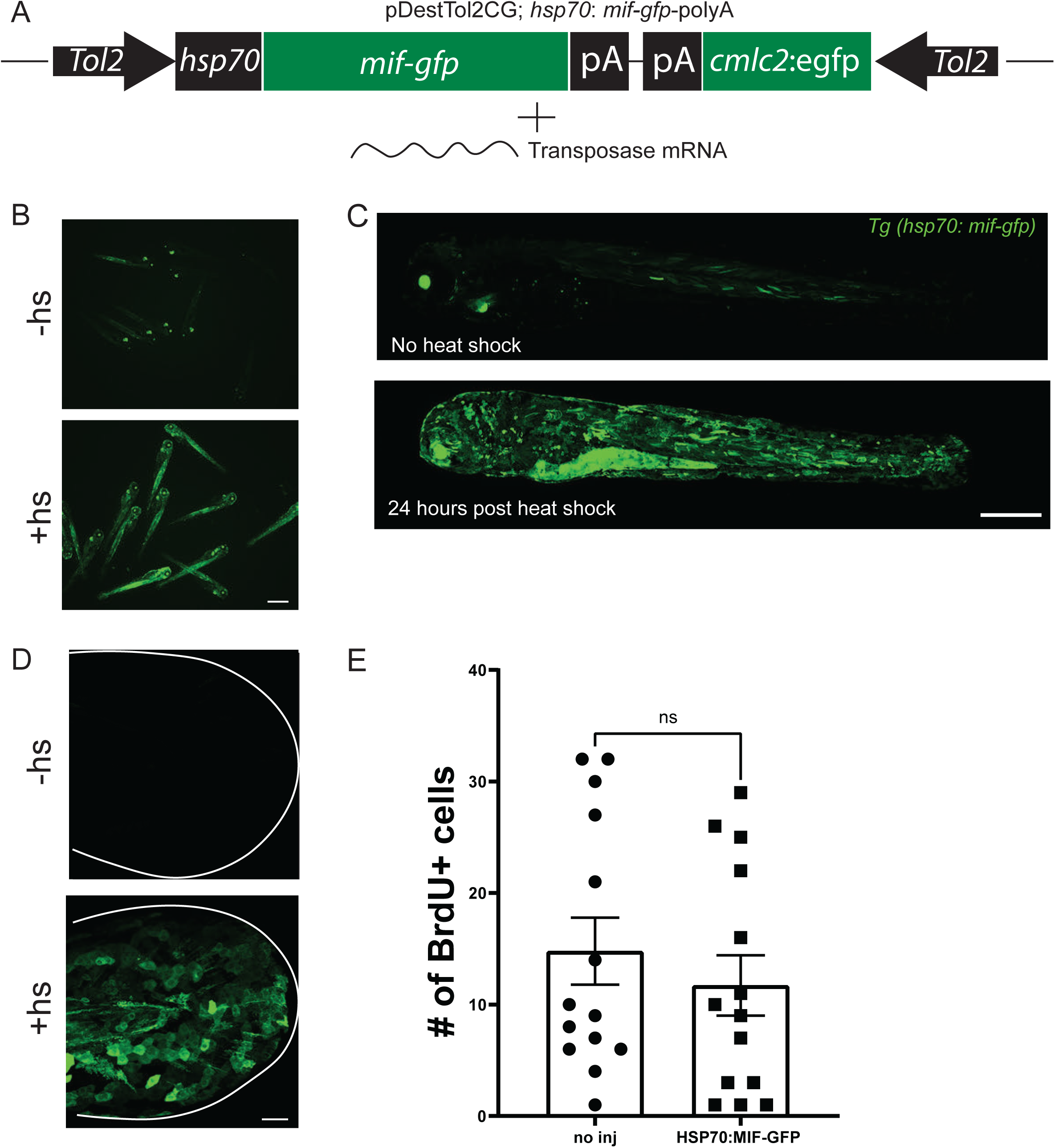
Heat-shock induction of MIF-GFP does not induce proliferation under homeostatic conditions. (A) Schema of the Tol2 construct used to drive *mif-gfp* downstream of the hsp70 promoter. Encoded within the genetic construct is a green heart marker using *cmlc2:gfp* to initially pick select larvae with the construct. All constructs were co-injected with transposase mRNA. **(B)** Representative large-field images of a clutch of zebrafish larvae before and after heat-shock induction of MIF-GFP. **(C)** A 10x confocal image of the distribution of MIF-GFP in a larvae pre and post heat shock induction. (D) A 20x confocal image of the variety of cell types within the tail epithelium expressing MIF-GFP. **(E)** A comparison of proliferation in heat-shocked and non-heat shocked larvae. n = 36, no inj. n = 32, heat shock. A student’s t-test was used to assess differences in means. Scale bars: B = 100 µm, C = 500 µm, D = 50 µm.

**Supplemental Figure 6.**
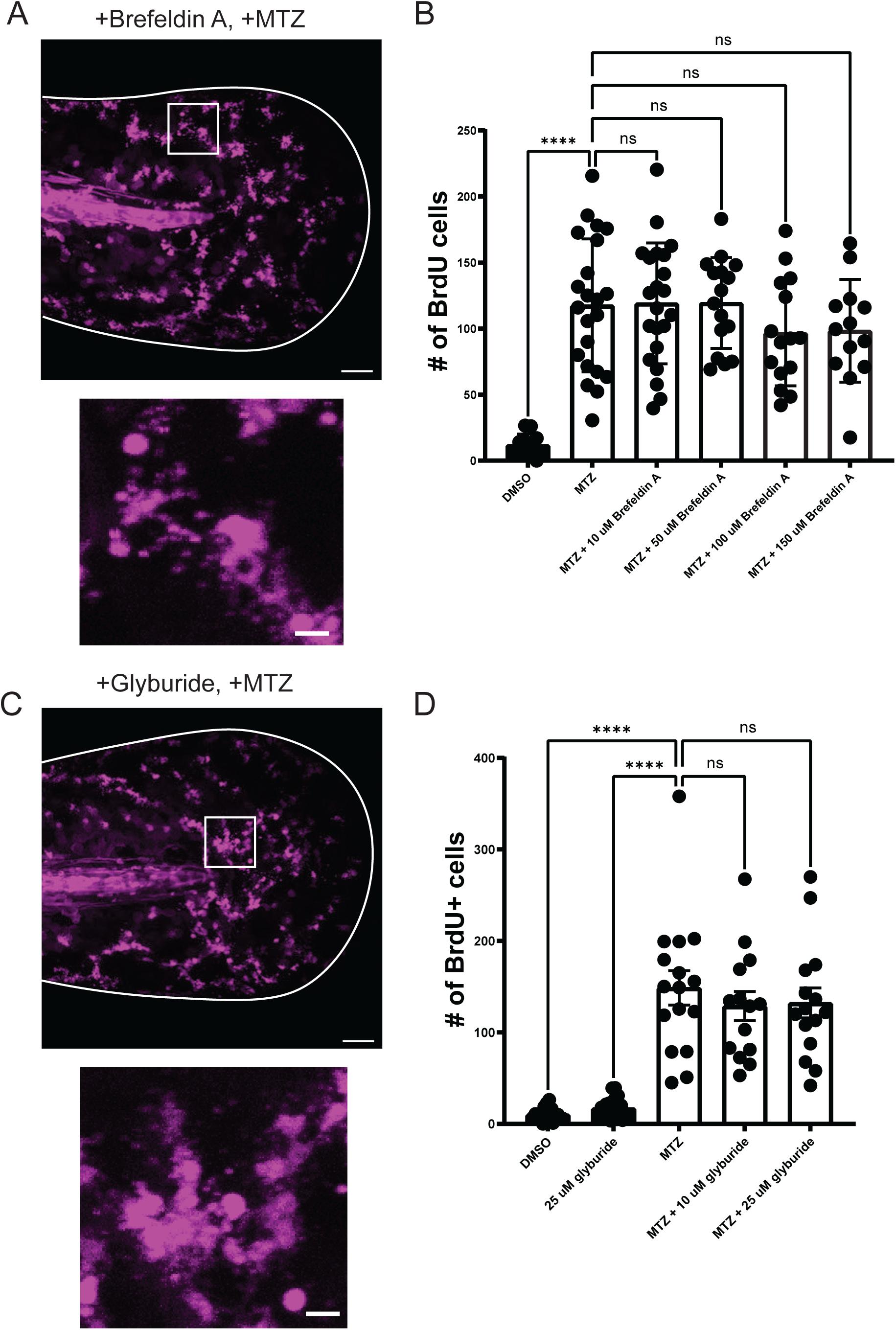
Treatment with secretion inhibitors do not alter apoptosis-induced proliferation. (A)Treatment with Brefeldin A does not affect AEV formation. (B) Treatment with various concentrations of Brefeldin A does not induce a significant reduction in proliferation. (C) esAEV formation after treatment with glyburide. (D) Apoptosis-induced proliferation in the presence of glyburide. Each condition had 32-40 larvae. Statistical significance was calculated with a Two-way ANOVA with a Tukey’s post hoc test. Scales bars = 50 µm.

**Supplemental Figure 7.**
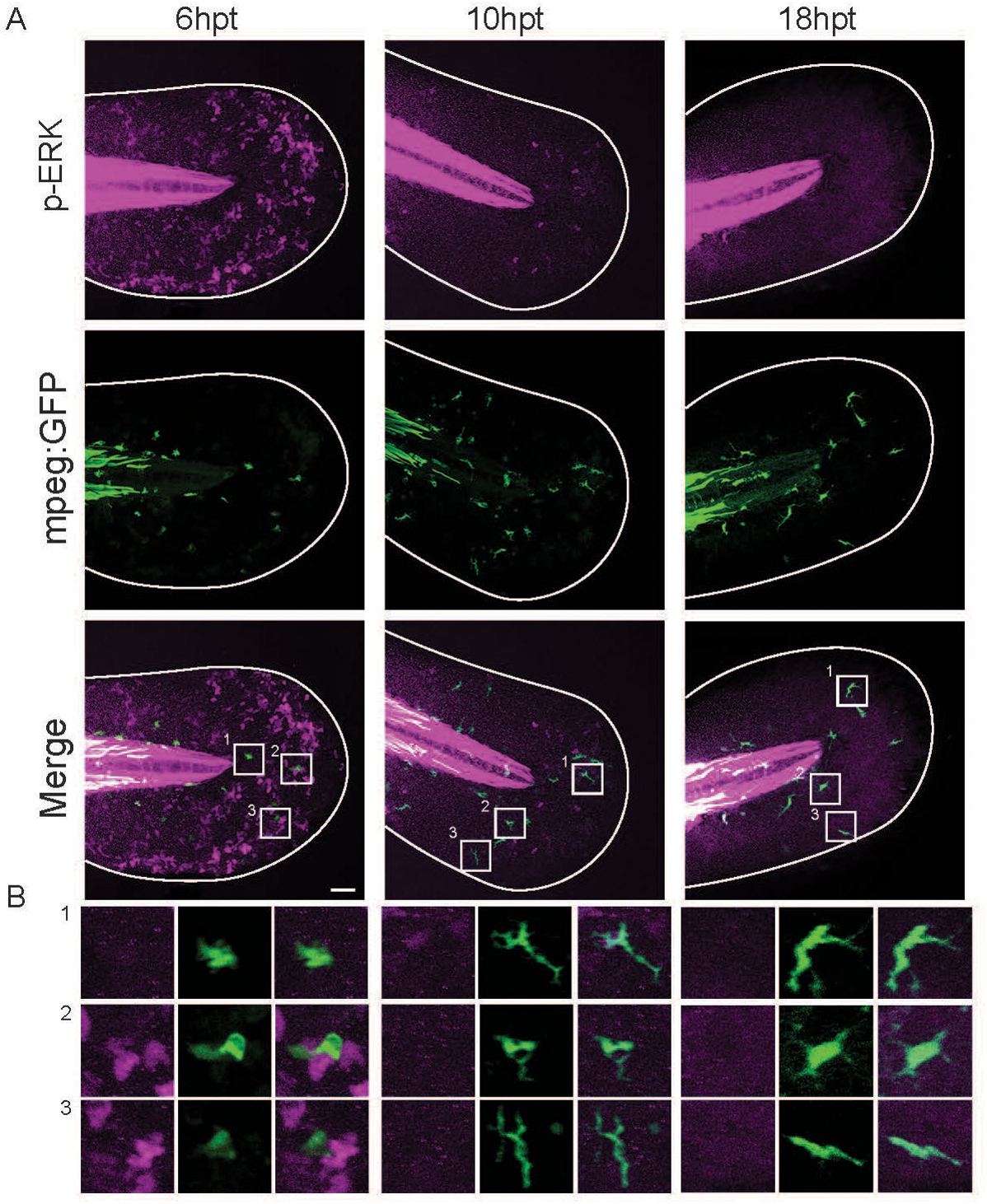
p-ERK signaling was not detectable in macrophages. (A) Representative images of p-ERK signaling in an *mpeg:GFP* background across three different timepoints post-MTZ treatment. (B) Images of 3 ROIS selected per timepoint highlighting the p-ERK level in macrophages. Scale bar = 50 µm.

**Supplemental Figure 8.**
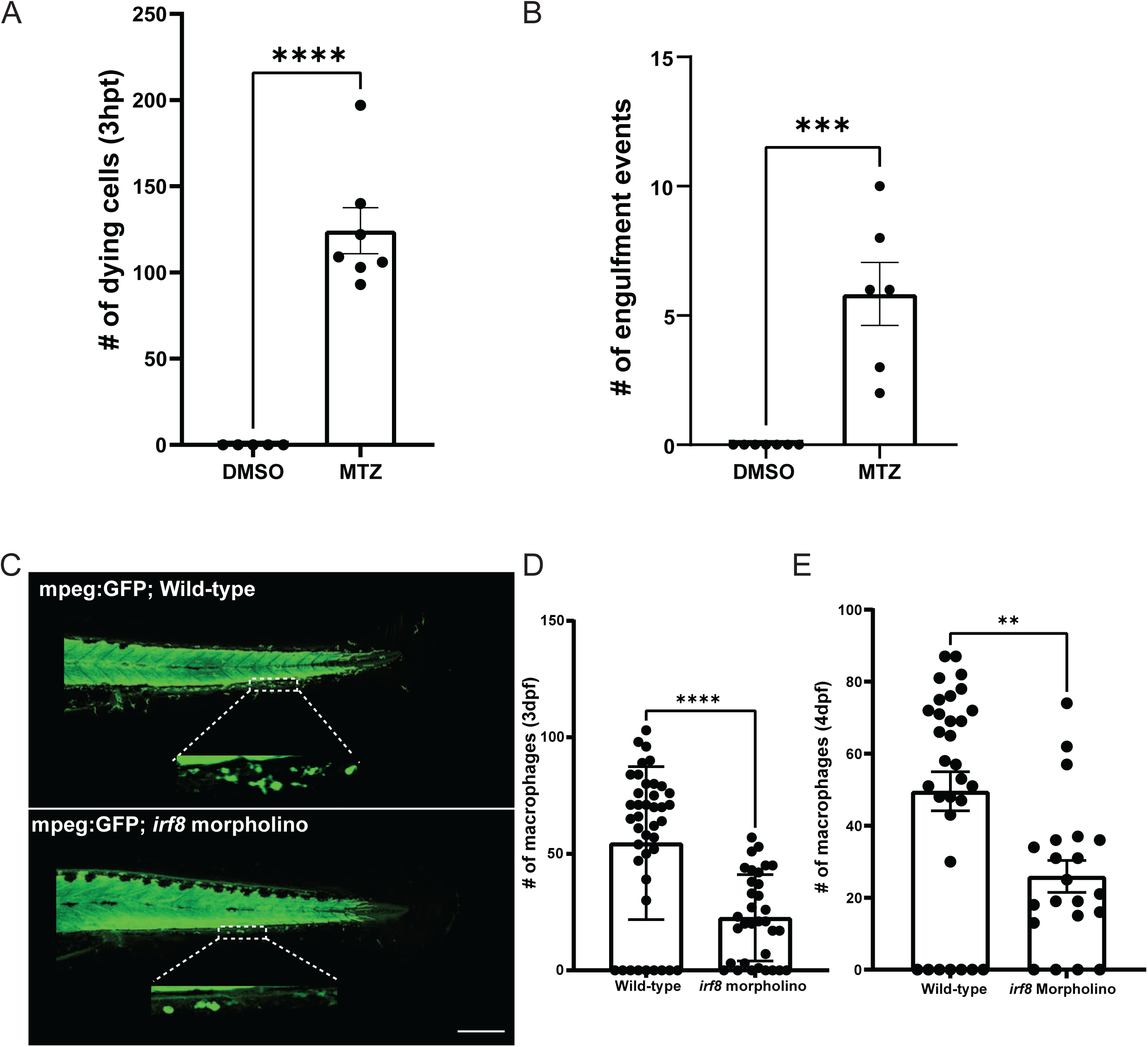
Macrophage contribution to AEV engulfment and depletion with *irf8* morpholino. (A)The number of epithelial stem cells that undergo apoptosis up to 3hrs post treatment. n = 5 larvae, DMSO. N = 6, MTZ. **** <0.0001 using a two-tailed t-test. (B) The number of engulfment events of macrophages in the presence of apoptotic cells across an 8-hour timespan. n = 8 larvae, DMSO. n = 6, MTZ. *** 0.0003 using a two-tailed t-test. (C) Ablation of the macrophage lineage using *irf8* morpholino (scale bar =200 µm). (D) Quantifications of the number of macrophages at 3dpf. n= 72 for Wild-type and n = 53 for *irf8* morpholino. ****<0.0001 via an unpaired two-tailed test. (E) Quantifications of the number of macrophages at 4dpf. n = 31, Wild-type. n = 21, *ifr8* morpholino. ** 0.0029 using an unpaired two-tailed t-test. Quantifications for D,E were performed by counting the number of macrophages present in the caudal vein from the cloaca to the tip of the notochord. Scale bar = 200 µm.

## SUPPLEMENTAL TABLES

Table 1. Proteins identified in the proteomic analysis of esAEVs.

## SUPPLEMENTAL MOVIES

**Supplemental Movie 1: The formation of esAEVs.** Time-lapse imaging of NTR-mCherry cells undergoing apoptosis and forming esAEVs *in vivo*. Images were acquired every 5 minutes across a z-stack and compiled as a maximum intensity projection, 2.5 fps. Scale bar = 10 µm. Time = hh:mm:ss.

**Supplemental Movie 2: Epithelial stem cell engulfment of apoptotic extracellular vesicles.** Time-lapse imaging of a p63:GFP positive cell engulfing an NTR-positive esAEV. Images were acquired every 15 minutes across a z-stack for 5 hours. 2.5 fps. Scale bar = 25 µm. Time = hh:mm:ss.

**Supplemental Movie 3: Macrophage dynamics during homeostasis and after induced apoptosis.** Time-lapse imaging of macrophage dynamics in an NTR-mCherry background. (Left) Macrophage movement with DMSO treatment. (Right) Macrophage movement after 4 hours of MTZ treatment. Images were acquired every 5 minutes across a z-stack for 8 hours. 5.0 fps. Scale bar = 50 µm. Time = hh:mm:ss.

